# Pairwise Attention: Leveraging Mass Differences to Enhance De Novo Sequencing of Mass Spectra

**DOI:** 10.1101/2025.03.28.645943

**Authors:** Joel Lapin, Alfred Nilsson, Mathias Wilhelm, Lukas Käll

**Author notes:** Equal contribution (Joel Lapin, Alfred Nilsson).

## Abstract

A fundamental challenge in mass spectrometry-based proteomics is determining which peptide generated a given MS2 spectrum. Peptide sequencing typically relies on matching spectra against a known sequence database, which in some applications is not available. Deep learning-based de novo sequencing can address this limitation by directly predicting peptide sequences from MS2 data. We have seen the application of the transformer architecture to de novo sequencing produce state-of-the-art results on the so-called nine-species benchmark. In this study, we propose an improved transformer encoder inspired by the heuristics used in the manual interpretation of spectra. We modify the attention mechanism with a learned bias based on pairwise mass differences, termed Pairwise Attention (PA). Adding PA improves average peptide precision at 100% coverage by 12.7% (5.9 percentage points) over our base transformer on the original nine-species benchmark. We have also achieved a 7.4% increase over the previously published model Casanovo. Our MS2 encoding strategy is largely orthogonal to other transformer-based models encoding MS2 spectra, enabling straightforward integration into existing deep-learning approaches. Our results show that integrating domain-specific knowledge into transformers boosts de novo sequencing performance.

## Introduction

Identifying peptide sequences from tandem mass spectrometry (MS2) is currently dominated by sequence searching, where spectra will be matched to in silico digests of sequences from a sequence database^1,2,3,4,5^. To obtain a high amount of identifications, one must choose a sequence database with a tenable search space size still containing sequences likely to be in the sample. Sequence searching inevitably has the weaknesses of bias and narrowness of the chosen sequence database, limiting the search only to those peptides the researcher believes will be present *a priori*.

De novo sequencing using deep learning^6,7,8,9^ (and traditional machine learning^10^) is an emerging approach that seeks to mitigate these weaknesses, wherein models can process the spectra and directly predict the peptide. These models, although not necessarily unbiased, can be trained on an expansive and diverse set of spectra, potentially overcoming the narrowness of sequence databases, provided that the model is reasonably accurate. Developed models have begun to be applied in areas where sequence searching is challenging. This can include applications where the search space is naturally large, such as immunopeptidomics^11^, or the uncertainty around the content of the analyzed sample makes choosing a reference sequence database very difficult, as in antibody-sequencing^12^, forensic samples^13^ and metaproteomics studies^7,8^.

The current state-of-the-art in de novo sequencing uses transformer models^14^ that frame de novo sequencing as a sequence-to-sequence translation problem^15^. In these models, a list of spectral peaks is encoded by transformers into a latent representation using self-attention. The latent representation is further processed by a decoder model, which predicts a matching peptide’s amino acid sequence in an autoregressive manner, using cross-attention. Human experts often interpret MS2 spectra by seeking common patterns of backbone cleavages, e.g. b- and y-ions, for successive peaks that differ by the mass of single amino acids or small fragments of a peptide^16^. Unlike human experts, deep learning models do not necessarily rely on predefined domain knowledge; instead, these models learn features by connecting the input to the target output using appropriate architectures, in our case the m/z and intensity peaks to a predicted amino acid sequence. A heavily parameterized model will automatically learn the features that best reduce the classification loss on the predicted sequence through gradient descent. However, such features are not easily decipherable, making these models practically uninterpretable, i.e., “black boxes”.

Despite the effectiveness of transformer models across a wide range of applications, leveraging domain knowledge can often improve performance beyond a direct naive implementation. In many applications, expert knowledge of the underlying problem is leveraged alongside the feature extraction capability of deep-learning modeling. In computer vision, convolutional neural networks^17^ incorporate an inductive bias by focusing on local receptive fields via learned filters and achieve state-of-the-art performance in many image tasks^18,19,20,21^. An especially informative example of domain knowledge alongside deep learning is AlphaFold2^22^ in protein structure prediction. AlphaFold’s Evoformer is a deep learning module that processes evolutionary information from multiple-sequence alignments, including the modeled sequence and amino acid pairwise structural features of a protein structure. This is an architecture specifically designed for protein structure prediction. The peculiarities of the AlphaFold2 model, and how they relate to its specific field/problem of protein structure prediction, suggest that performance improvements can be achieved through careful consideration of domain-specific data and mechanisms underlying the problem.

Herein, we report the improvement of de novo sequencing by a transformer model incorporating features inspired by the traditional ways human experts would annotate an MS2 spectrum. We term our variation on the transformer as the Pairwise Attention model (PA), which concentrates on modifying the encoder half of the transformer to optimally process spectra into a latent space, used then for decoding the sequence. Specifically, for a spectrum of length N, we create an NxN set of features whose entries are the m/z differences of all pairs of peaks in the spectrum. Inspired by AlphaFold2’s pair representations of protein sequences, we feed this antisymmetric pairwise matrix into the transformer blocks as an attention bias. This augmentation to the transformer architecture is lightweight and programmatically simple to implement. We see large improvements over our own implementation of a transformer without such pairwise features, achieving a 14.2% (6.7 percentage point) increase in average peptide precision at 100% coverage over our base model and a 7.2% (3.6 percentage point increase) over Casanovo, when tested on the revised nine-species benchmark.

## Methods

Our model follows a standard encoder-decoder transformer architecture, but we modified the encoder’s self-attention mechanism to incorporate pairwise m/z differences as an additive bias.

### Peak embeddings

The sequence of *N* peaks, each consisting of a mass and intensity 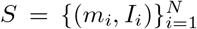, is fed into the transformer blocks per the standard spectrum encoding scheme, following PointNovo^6^ and Casanovo^7^. Specifically, the *N* peaks of m/z and intensity values are expanded into Fourier features of dimension *r*_*m*_ and *r*_*I*_, respectively. Equations (1a) and (1b) represent the processing of the original spectrum.

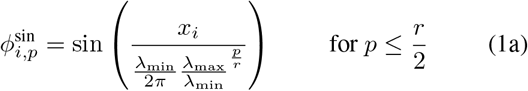

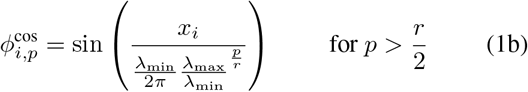

where *x*_*i*_ could either be the mass or the intensity of the *i*-th peak. Here, *p* indexes the Fourier feature dimension, and *λ*_max_ and *λ*_min_ are hyperparameters controlling the wavelength range.

The resulting Fourier features for each peak are concatenated along the feature dimension, producing an intermediate matrix of shape *N ×* (*r*_*m*_ + *r*_*I*_). To transform this into the final input token matrix for the transformer, a linear projection is applied:

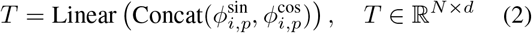

where *d* is the desired input feature dimension for the transformer encoder.

### Pairwise features

The 2D pairwise features (two sequence dimensions) are constructed by computing the pairwise differences between all pairs of peaks in the input spectrum. Specifically, for a sequence of *N* peaks, the pairwise m/z difference matrix Δ*m* ∈ ℝ^*N ×N*^ is defined as:

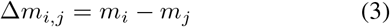

where *m*_*i*_ and *m*_*j*_ are the m/z values of peaks *i* and *j*, respectively. To encode these pairwise differences, we expand each 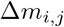 into Fourier features of dimension *r*_*pw*_*′* :

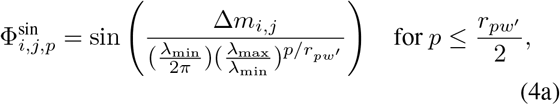

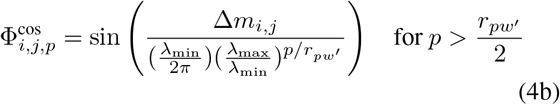

Let 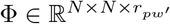 denote the resulting pairwise feature matrix.

### Pairwise attention

Our mechanism for pairwise attention borrows concepts from AlphaFold2’s “MSA row-wise gated self attention with pair bias” module, which must combine data of different modalities: a multiple-sequence alignment and pairwise amino acid encodings. Specifically, it feeds a pairwise representation of the sequence’s amino acids as a bias to the self-attention mechanism where keys, queries, and values originate from the multiple-sequence alignment. Our model mostly adheres to AlphaFold’s mechanism, but instead for Fourier features of the m/z sequence and for Fourier features of the pairwise m/z differences.

Recall that transformer self-attention computes attention weights between all pairs of input positions. Given input embeddings *X* ∈ ℝ^*N ×d*^ for *N* peaks, we first compute queries *Q*, keys *K*, and values *V* for each attention head *h* and encoder layer *l*:

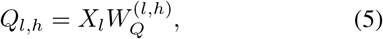

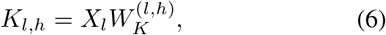

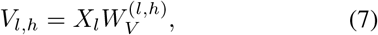

where 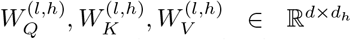 are learnable weight matrices, and *d*_*h*_ is the dimensionality per head, with *H* being the total number of heads.

The standard self-attention weights *A*_*l,h*_ for head *h* at layer *l* are then computed as:

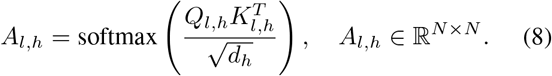

To incorporate pairwise m/z differences, we define a learned function *f*_pw_(Φ), of the pairwise feature matrix (Equation (4)). This function maps the pairwise features to a latent space with dimensionality 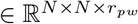. We then apply a *layer-specific* linear transformation *g*^(*l*)^, which adapts the pairwise bias across the network depth. This results in the following attention activation map for each head *h* and layer *l*.

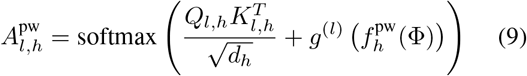

where 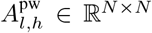. For 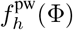, we use two linear transformations with a SiLU activation in-between:

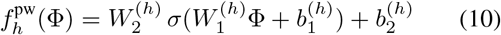

where 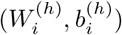 for *i* = 1, 2 are learnable parameters, shared across encoder layers, and *σ* is the SiLU activation function.

Each encoder attention module *l* applies its own linear transformation *g*^(*l*)^ to the output of *f* ^pw^:

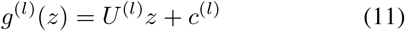

where 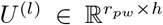 and *c*^(*l*)^ ∈ R^*h*^ are layer-specific parameters. This setup allows the network to adapt the pairwise bias across its depth.

### Memory footprint and computational cost

The pairwise features, each of which has dimension *r*_pw_*′*, are first linearly transformed to *r*_pw_ units through (*W*_1_, *b*_1_) of Equation 10. When training in batches this operation creates a matrix of size batch size × *N* × *N × r*_pw_, which can result in substantial memory overhead when processing spectra with a large number of peaks. For this reason, it is important to select a conservative value for *r*_pw_, typically smaller than *r*_*m*_ or *r*_*I*_.

Inside the self-attention module, the pairwise features are linearly transformed to have the same number of channels as there are attention heads (*h*), and then added as the attention bias before taking the softmax, over the keys dimension. An illustration of the PA mechanism is depicted in Figure S1. By this mechanism, the pairwise features can exert great influence on the resultant attention map after the softmax is taken.

The total number of parameters added by the pairwise features is rather insignificant compared to the rest of the transformer. The first transformation of the pairwise Fourier features (*W*_1_, *b*_1_) adds 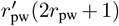 parameters, and (*W*_2_, *b*_2_) adds *r*_pw_(2*r*_pw_ + 1) parameters. *g*^(*l*)^ is a linear transformation for each self-attention module along the depth of the encoder. Each attention transformation has size *h*(*r*_pw_ + 1) parameters. For a depth of D, this is a total parameter count for the entire network of 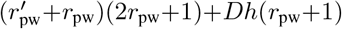. In this work, we use a model with 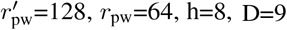 (see implementation details), which adds a total of 29,448 parameters, or about +0.1%, to the model. The VRAM memory cost increases from 9.3 GB (Base) to 17 GB (PA) for a single batch of 100 spectra.

### Data

We used two versions of the nine-species benchmark for comparison to other works. Our initial tests were on the original nine-species dataset, referred to as nine-species V1. This dataset is the smaller of the two, consisting of 1,526,282 total spectra. We downloaded Casanovo’s deposited preprocessed version of nine-species V1^9^, published on Zenodo in 2022 (https://zenodo.org/records/6791263). The nine-species version 2 (V2) dataset was re-searched by the Casanovo team to improve data quality and PSM confidence. This dataset contained 2,844,842 total spectra. At the time of this writing, this dataset was available in the Massive repository (ftp://massive.ucsd.edu/v05/MSV000090982/updates/2024-05-14_woutb_71950b89/peak/9speciesbenchmark/), where all relevant mgf files for each species can be downloaded and processed. For each version of the nine-species dataset, we parsed all modified sequences to enumerate the tokens and establish the token dictionary that the model would use when run on that respective dataset. The distribution of the number of spectra and peptides for each of the species in the set can be found in Table 1 To download the MassIVE-KB dataset, we downloaded a metadata file provided from the MassIVE-KB website (https://massive.ucsd.edu/ProteoSAFe/static/massive-kb-libraries.jsp).^23^ The metadata file provides filenames and URL links for mzML files, from which we obtained the matching PSMs in the dataset. We used 98.75% of the data for training, with a small development split of 1% for validation and 0.25% for testing. The true evaluation was done on each species of the nine-species dataset.

**Table 1.** The number of spectra and peptides in the nine-species dataset.

| SPECIES | SPECTRA COUNT<br>NINE-SPECIES V1 | UNIQ. PEP. NINE-<br>SPECIES V1 | SPECTRA COUNT<br>NINE-SPECIES V2 | UNIQ. PEP. NINE-<br>SPECIES V2 |
| --- | --- | --- | --- | --- |
| *Apis Mellifera* | 313,844 | 44,840 | 194,218 | 30,337 |
| *Bacillus Subtilis* | 291,769 | 34,258 | 1,358,337 | 66,455 |
| *Candidatus Endoloripes* | 150,117 | 16,959 | 82,290 | 10,884 |
| *Homo Sapiens* | 130,049 | 28,954 | 44,604 | 13,930 |
| *Methanosarcina Mazei* | 164,412 | 36,207 | 267,332 | 23,860 |
| *Mus Musculus* | 37,019 | 9,463 | 25,541 | 6,871 |
| *Solanum Lycopersicum* | 290,000 | 92,934 | 178,413 | 62,467 |
| *Saccharomyces Cerevisiae* | 111,298 | 30,174 | 585,593 | 33,199 |
| *Vigna Mungo* | 37,774 | 4,395 | 108,514 | 13,857 |
| TOTAL | 1,526,282 | 298,184 | 2,844,842 | 261,860 |

In addition to the nine-species benchmark, we evaluated our models on an independent bacterial dataset^24^ (PRIDE accession number PXD010613). This dataset contains only variable modification of oxidized methionine and no fixed modifications. Because this dataset is of bacterial origin, it is distinct from the MassIVE-KB training data, and the possibility of peptide leakage into the test set should be substantially lower than that for nine-species, providing a more rigorous test of model generalization.

### Adjustment of PEPMASS annotation

During data preprocessing, we identified inconsistencies in the `PEPMASS` annotations within the `mgf` files of both nine-species V1 and V2 datasets. Specifically, the `PEPMASS` field erroneously contained the precursor m/z values instead of the precursor masses. In contrast, the MassIVE-KB dataset’s `PEPMASS` field contained the precursor mass calculated as the product of the precursor m/z and charge, but without accounting for the mass of the protons (i.e., it did not subtract the proton masses associated with the charge state). It is worth acknowledging that there is no official standard for this field - the community does not agree upon the content, but according to MASCOT documentation^1^, it should be populated with the peptide mass.

To ensure consistency across datasets, we addressed these discrepancies by adjusting the `PEPMASS` values in the nine-species V1 and V2 datasets. Specifically, we multiply the precursor m/z value in the `PEPMASS` field by the charge state and set this product as the new `PEPMASS` value:

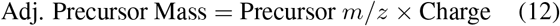

This adjustment aligns the `PEPMASS` annotations in the nine-species datasets with those in the MassIVE-KB dataset, where the `PEPMASS` field already contains this product.

While this correction does not account for the total proton mass associated with the charge state (i.e., it does not subtract Charge × *m*_*p*_, where *m*_*p*_ = 1.00727647 Da is the proton mass), incorporating the proton mass offset is unlikely to affect our machine learning model’s performance.

### Implementation details

In order to have a fair comparison we largely adhered to Casanovo’s model configuration and hyperparameter settings for the PA model. We trained with 150 top intense peaks in a spectrum, and a maximum peptide length of 100. Spectral intensities were divided by the base peak, such that the base peak has *I*_*max*_=1 and all other peaks are scaled downward. The architectural parameters chosen for the spectral features from Eq. 1 were the following: 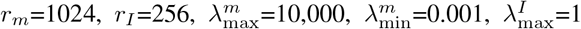and 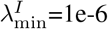. The parameters chosen for the pairwise features from Eq. 4b were *r*_*pw*_*′* = 128, and the same frequencies as the spectral features. After concatenation of intensity and mass Fourier features, the tensor is linearly projected back to *r* = 512 units. The pairwise features are projected back to *r*_*pw*_ = 64; a smaller value than the 1d features was chosen to help stay within our GPU’s memory budget. Attention modules in the encoder were constructed with *d* = 64 query, key, and values units, *h* = 8 attention heads and 9 total attention blocks. The encoder was a custom implementation in Pytorch and had a total of 19.6 million parameters. The decoder had all the same hyperparameter settings as the encoder, but had 28.4 million parameters due to the extra cross attention modules. The decoder was implemented through the Depthcharge^25^ library (the same used by Casanovo) and altered to be compatible with our specific processing of the data. Dropout on both the encoder and decoder was set to 0.25.

We trained the PA model using the cross entropy loss function, implemented with Teacher forcing, where an attention mask prevents any token in the decoder from attending to future positions. Models were trained for 50 and 40 total epochs, for nine-species V1 and V2, respectively. A batch size of 100 was used. The learning rate was linearly warmed up to 2e-4 in 20,000 steps, and then held constant for the remainder of training. We used the Adam optimizer with default PyTorch parameters.

For the MassIVE-KB dataset, instead of matching hyperparameters for comparison’s sake, we sought to train a more optimal realization of our model. We trained models that included 300 top intense peaks and tested on the nine-species V2 dataset with a 5-beam beam search. Here, the learning rate was linearly warmed up to 2e-4 in 600,000 steps, and then held constant for the remainder of training. Models were trained for 10 epochs, after which the checkpoint with the best validation score was selected for final testing.

### Evaluation metrics

All results for nine-species follow the procedure of training on 8 species and testing on the holdout 9th species. We report precision at 100% coverage (no confidence cutoff) at the peptide level. This is calculated as the number of predicted sequences that match the ground truth peptide divided by all peptides in the testing species’ dataset, *N*_match_*/N*_total_. We follow the amino acid matching methodology first introduced by DeepNovo,^26^ wherein matched amino acids must be *<* 0.1 Da and have a prefix or suffix mass that differs from the ground truth by *<* 0.5 Da. To ensure that we don’t simply optimize a specific random seed and have a robust result, all reported precision values for PA are the average of 3 random seeds, 0, 10, and 20. We only compare to Casanovo’s reported statistics, specifically those without a beam search. Casanovo has multiple published preprints with varying numbers for the nine-species benchmark; we specifically compared our results to their reported results in their most recently published article^7^, provided in their supplementary materials.

For calculating the peptide confidence for the precision-coverage curves, we closely followed Casanovo’s peptide score metric. We take the mean softmax confidence of the predicted amino acids up to the stop token and set the confidence to ™ 1 for any predicted peptide’s mass that was more than 50 ppm off the precursor mass.

## Code availability

All code is available for implementation and reproducibility of our work at https://github.com/statisticalbiotechnology/pairwise.

## Results

### Nine-species benchmark

We assess the PA model’s performance on the nine-species benchmark in two ways: 1) our results for our encoder with pairwise attention (PA), against the same model without pairwise attention, which we refer to as Base, and 2) our implementations against the reported results for Casanovo_*bm*_, which is their implementation without a beam search^7^. Casanovo reported improvements over their top predecessors, PointNovo^6^, DeepNovo^9^, and Novor^10^.

The results and model comparisons for nine-species V1 are displayed in Figure 2 as peptide precision/coverage curves for all nine-species and the comparisons of precision at 100% coverage in Table 2. Our PA model was run at 3 different seeds, 0, 10, and 20, and is reported with its standard deviation. When looking at average precision over all nine-species in Table 2, PA gives a boost of 11.3% (5.9 percentage points) peptide precision over the Base model. This is a substantial improvement given that PA adds only 29.5k more parameters, or approximately a 0.1 percentage points increase in the encoder size. Amongst the nine-species, the increases in performance range from 4-6 percentage points (*Candidatus Endoloripes, Homo Sapiens, Methanosarcina Mazei, Mus Musculus*, and *Vigna Mungo*) to over 7 percentage points (*Apis Mellifera, Bacillus Subtilis, Saccharomyces Cerevisiae*, and *Solanum Lycopersicum*).

**Table 2.** Peptide precision at 100% coverage as measured on the V1 and V2 of the nine-species dataset. Here, Base denotes our implementation of a standard transformer, i.e. without pairwise features. PA is our implementation of pairwise attention, and Casanovo is the reported numbers for Casanovo_bm_ in the original publication^7^. For PA we report the average of 3 runs at seeds 0, 10, and 20, with standard deviation in parentheses.

| TEST SET | NINE-SPECIES V1 |  |  | NINE-SPECIES V2 |  |  |
| --- | --- | --- | --- | --- | --- | --- |
| MODEL | BASE | PA | CASANOVO | BASE | PA | CASANOVO |
| *Apis Mellifera* | 0.390 | **0.463 (0.004)** | 0.433 | 0.390 | 0.446 (0.009) | **0.456** |
| *Bacillus Subtilis* | 0.536 | **0.612 (0.009)** | 0.573 | 0.494 | **0.583 (0.006)** | 0.538 |
| *Candidatus Endoloripes* | 0.357 | **0.409 (0.002)** | 0.390 | 0.356 | 0.425 (0.006) | **0.468** |
| *Homo Sapiens* | 0.340 | **0.391 (0.003)** | 0.383 | 0.451 | 0.521 (0.004) | **0.533** |
| *Methanosarcina Mazei* | 0.503 | **0.554 (0.002)** | 0.515 | 0.509 | **0.579 (0.012)** | 0.529 |
| *Mus Musculus* | 0.433 | **0.472 (0.003)** | 0.431 | 0.410 | **0.435 (0.004)** | 0.395 |
| *Solanum Lycopersicum* | 0.509 | **0.590 (0.008)** | 0.522 | 0.543 | **0.623 (0.007)** | 0.608 |
| *Saccharomyces Cerevisiae* | 0.537 | **0.612 (0.009)** | 0.580 | 0.558 | **0.631 (0.010)** | 0.561 |
| *Vigna Mungo* | 0.570 | **0.625 (0.002)** | 0.552 | 0.525 | **0.598 (0.022)** | 0.428 |
| AVERAGE | 0.464 | **0.523 (0.004)** | 0.487 | 0.471 | **0.538 (0.009)** | 0.502 |

**Figure 1.**
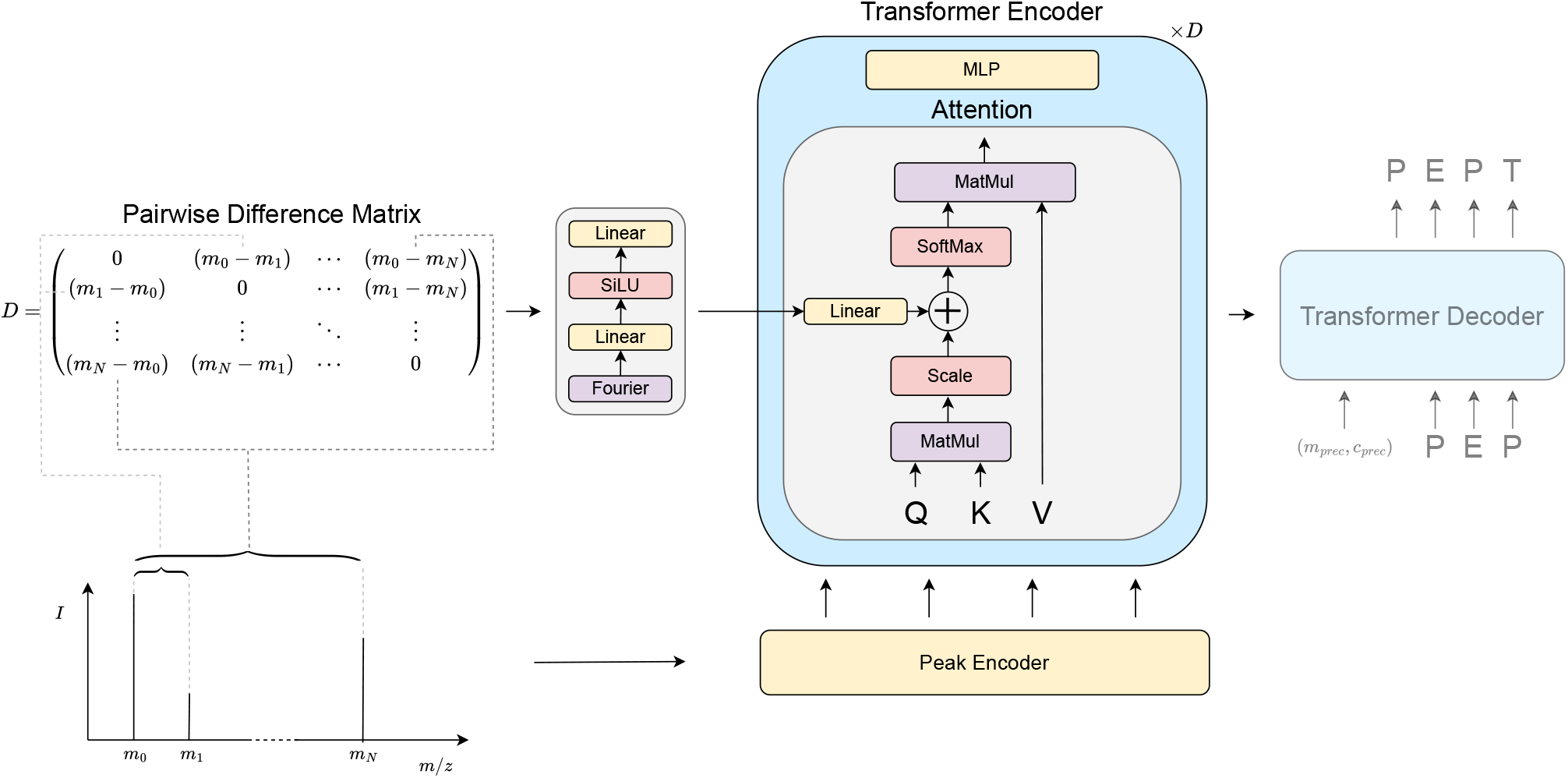
Architecture of the Pairwise Attention model, depicted through the self-attention mechanism of the Transformer encoder. The original mass spectrum in the lower left is turned into 1D features via the peak encoder, which concatenates fourier features of the m/z and intensity dimensions, and further processed into 2D features by taking the pairwise differences of its m/z values. As the 1D features are processed by standard Transformer encoder modules, i.e. self-attention and multi-layer perceptron (MLP) networks, the 2D features are fed into the self-attention module as a learnable bias before the softmax attention. This bias is fed into self-attention mechanisms throughout the depth of the encoder. The Transformer decoder is unaltered from the original implementation.^7, 14^

**Figure 2.**
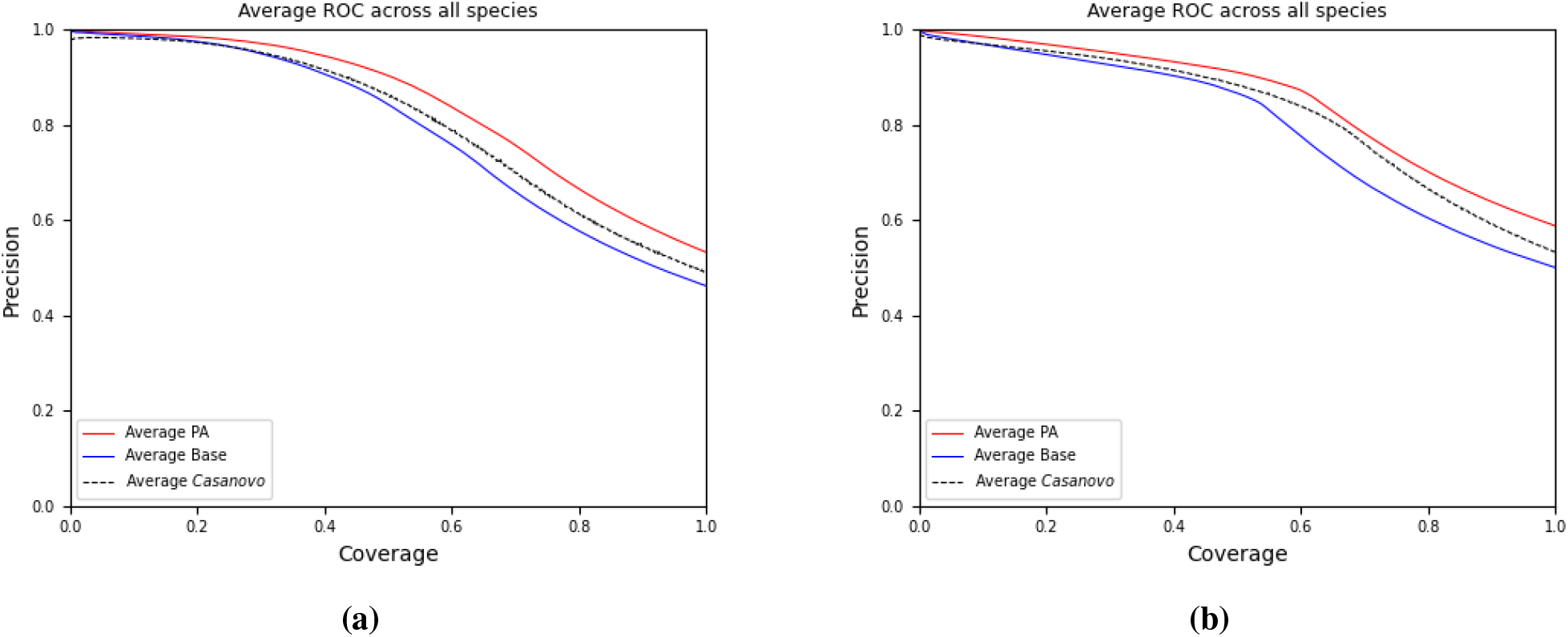
Precision-coverage curves for our PA and base models, and Casanovo’s reported BM model. Nine-species V1 is displayed in a) and V2 is displayed in b). For the PA model, only the best of the 3 seeds is plotted.

When compared to the published results on the V1 dataset, the PA model improves on Casanovo by 7.4% (3.6 percentage points). PA beats Casanovo for every species, with an exceptional increase of 6.8 percentage points for *Solanum Lycopersicum*. It is important to make note of the disparity between our Base transformer implementation and Casanovo’s reported numbers, which is a considerable 2.3 percentage points on average. Non-optimal hyperparameters may partly explain worse performance than Casanovo’s reported results, but is not sufficient to fully account for the overall difference in performance. It is possible that subtle different implementations of training/evaluation code and processing of data explain the inability to reproduce Casanovo’s original result, but were ultimately not identified in this study.

The nine-species V2 dataset, constructed by the Casanovo authors, is reported to be higher confidence PSMs for each species. As of the time of this writing, Casanovo is the only other published model we are aware of that has reported results for this dataset. When compared to the Base architecture, we see a very consistent increase in performance for PA overall, similar to the V1 dataset. The average peptide precision increases by 14.2% (6.7 percentage points). A modest difference from V1 to V2 is not unexpected, as the updates to the nine-species dataset significantly changed the size and likely also the quality of the data, but the persistence of the trends in improvement for all species shows that the advantage from pairwise attention is a robust and consistent result amongst our implementations.

When compared to Casanovo, again we see an improvement for our PA model by 7.2% (once again by 3.6 percentage points) in average peptide precision at 100% coverage. One important observation to make is how much more variability there is in the comparison of the two models than with nine-species V1. In nine-species V1, PA was consistently above or equal to Casanovo for all species. This is evident in the precision-coverage curve of Figure 2a, where PA’s precision lies mostly above Casanovo’s precision for all confidences, for all V1 species. For nine-species V2 in Figure 2b Casanovo has equal or higher precision, over various confidence ranges, for *Apis Mellifera, Candidatus Endoloripes, Homo Sapiens*, and *Solanum Lycopersicum*. When looking at performance at 100% confidence, in Table 2, the increases for our PA model range from 0-6 percentage points. Our model is performing worse for 3 species, *Apis Mellifera, Candidatus Endoloripes*, and *Homo Sapiens*; Casanovo exceptionally had 4.3 percentage points better peptide precision for *Candidatus Endoloripes*, whereas PA was better than Casanovo for this species by 1.9 percentage points on V1. For the rest of the species, our model was better, especially for *Vigna Mungo*, which was 17 percentage points better than Casanovo while only 7 percentage points better on V1. Changes in the quality of the data notwithstanding, this result seems questionable because of the inconsistency with the results of V1 and the two models’ seeming unusual ability to specialize on specific train/test data splits. Furthermore, in contrast to Casanovo, the comparison between our Base and PA models is consistent across the two datasets. To reconcile this apparent incongruence in the results, it would be best to faithfully implement all models within the same platform/code base. We reserve this comparison for future work.

As a final demonstration on nine-species, we trained both our Pairwise Attention (PA) and Base models on the MassIVE-KB dataset, using beam search and an increased maximum number of peaks per spectrum. The best-performing model was then evaluated on all species in the nine-species V2 dataset; the results shown in Table 3. Consistent with previously reported results for Casanovo^7^, training on the MassIVE-KB dataset led to substantial performance improvements compared to training on the smaller, lower-quality nine-species dataset. The improvement of PA over Base, as shown in Table 3, is now 1.6% (2 percentage points on average) - narrower than when training on the V2 dataset alone. This narrowing is expected with larger training datasets. However, the persistence of a clear gap between PA and Base, even with such an extensive dataset (30M spectra), underscores the significance of our results and suggests that the gains from our learned pairwise attention bias cannot easily be diminished by further scaling the training data.

**Table 3.** Performance after training the models on the MassIVE-KB set. Here we tested on each of the species in the nine-species dataset and report peptide precision at 100% coverage. The models tested are Base and PA, alongside Casanovo’s reported numbers^7^.

| TEST SET | NINE-SPECIES V2 |  |  |
| --- | --- | --- | --- |
| MODEL | BASE | PA | CASANOVO |
| *Apis Mellifera* | 0.618 | 0.640 | **0.662** |
| *Bacillus Subtilis* | 0.706 | 0.732 | **0.778** |
| *Candidatus Endoloripes* | 0.525 | 0.549 | **0.656** |
| *Homo Sapiens* | 0.737 | **0.746** | 0.740 |
| *Methanosarcina Mazei* | 0.700 | **0.720** | 0.710 |
| *Mus Musculus* | 0.563 | **0.579** | 0.552 |
| *Solanum Lycopersicum* | 0.735 | 0.751 | **0.799** |
| *Saccharomyces Cerevisiae* | 0.765 | 0.784 | **0.840** |
| *Vigna Mungo* | 0.753 | **0.783** | 0.762 |
| AVERAGE | 0.678 | 0.698 | **0.722** |

When compared to Casanovo, the PA model improved for *Homo Sapiens, Methanosarcina Mazei, Mus Musculus*, and *Vigna Mungo*, and trails Casanovo for *Apis Mellifera, Candidatus Endoloripes, Solanum Lycopersicum*, and *Saccharomyces Cerevisiae*. Notable amongst these latter species is the extent to which Casanova outperforms our PA model, by ∼ 5 percentage points for *Solanum Lycopersicum* and *Saccharomyces Cerevisiae*, and exceptionally by 10.7 percentage points for *Candidatus Endoloripes*.

### External bacterial dataset

Although the nine-species benchmark is the most widely used dataset for de novo sequencing studies, its limitations - including the age and quality of the spectra and instruments used for collection - raise concerns about its suitability for comparing deep learning models. While it remains useful for model prototyping, a performance comparison on a more contemporary dataset is desirable. To address this, we evaluated our MassIVE-KB-trained models (without a beam search) on an independent bacterial dataset and compared its performance to Casanovo’s own publicly available model checkpoint and code. Additionally, because the external bacterial dataset is vastly different in origin from the MassIVE-KB training set, the likelihood of peptide leakage is minimal, making it a better test of generalization. Through this evaluation, we can allay concerns about the idiosyncrasies of the nine-species benchmark.

Table 4 reveals that the PA model lies 2.5 percentage points above Base, consistent with MassIVE-KB results in Table 3, but now both PA and Base considerably improve over Casanovo. On the bacterial dataset PA is 8.5 percentage points (24% relative improvement) above Casanovo, in great contrast to the difference observed between the same model checkpoints evaluated on nine-species V2 and Casanovo’s reported numbers. Even more striking is that Base is now well above Casanovo’s performance, which was not the case in either Table 3 or the nine-species cross-validation in Table 2. It is important to emphasize that no hyperparameter tuning was done for training our models (or Casanovo) on this dataset, and PA consistently improves over the Base model. As stated above, it is a future priority when comparing models to always run on the same data, in the same platforms, for fair comparison, as demonstrated here on the bacterial dataset.

**Table 4.** Performance on an external bacterial dataset after training the models on the MassIVE-KB set. Here we tested on the bacterial test dataset and report peptide precision at 100% coverage. The models tested are Base, PA, and Casanovo’s own hosted MassIVE-KB model checkpoint and code^7^.

| MODEL | PRECISION |
| --- | --- |
| BASE | 0.408 |
| PA | **0.433** |
| CASANOVO | 0.348 |

### Runtime cost

While the inclusion of pairwise features comes at a very light parameter cost, the increase in feature maps with sequence size (*N*, top peaks) that must be produced (and saved for backpropagation) by the model slows down forward and backward passes. For 3 runs of nine-species V1 with *Vigna Mungo* as the holdout species, which was one of the species with the fewest spectra and thus one of the largest training sets, the average total training time was 90,177 seconds for 446,467 total training steps, or 4.95 training steps per second. For the Base encoder the average total training time was 62,291 seconds, 7.17 training steps per second, which is 31% faster. The cost in runtime must be weighed by both developers and users of the model, depending on application. All nine-species models were trained on a single A100 GPU, and the MassIVE-KB models on 16 A100 GPUs.

## Discussion

The results provided here show a substantial improvement in de novo sequencing performance when incorporating features that encode pairwise differences between all peaks in a spectrum. Specifically, our model with pairwise features demonstrates a substantial improvement over our baseline model without these features and a modest improvement over published results on the nine-species benchmark. The idea of adding such features was inspired by the intuition used when manually annotating spectra, where one often identifies sequences from pairs of peaks that differ by the masses of modified and unmodified amino acids.

It can be argued that handcrafted features or inductive biases are unnecessary, as transformers have the capacity to learn higher-ordered features such as the pairwise differences and their correspondence to different fragments on their own. We see evidence suggesting that during training, the Base model may be learning representations of pairwise features, or approximations of them. For example, the pairwise model consistently shows a faster decrease in training loss and improved validation peptide precision compared to the base model (Figure S2), although the performance gap narrows over time. Nevertheless, after both models converge to their best validation scores, the pairwise model continues to outperform the Base model in the validation metrics and ultimate test performance.

This suggests that the transformer’s capacity to solve de novo sequencing may be limited when relying on randomly initialized models optimized by gradient descent. Handcrafted features, such as the pairwise distances, can direct the model to converge faster. It is also possible that the inclusion of high order features as input from the start of training might relieve the model to learn such features from scratch allowing the procedure to focus on the learning of even higher ordered features. Our results show that feature engineering, which traditionally has been essential for machine learning techniques such as decision trees and support vector machines, can still play a valuable role in deep learning alongside raw inputs.

The PA model was better than the Base model for both the nine-species dataset V1 and V2, but then only slightly better for each respective architecture trained on MassIVE-KB. It is possible that pairwise attention is most beneficial when training data is limited, since MassIVE-KB is more than 10 times the size of nine-species V2. Other possibilities are that PA is most helpful when data quality is low and that MassIVE-KB contains higher-quality PSMs than the nine-species dataset. Further testing is needed to elucidate the advantages of PA, and in what situations they may be marginalized.

It should be noted that our model has the same transformer implementation of the peptide decoder as Casanovo (Depthcharge), with architectural differences only in the encoder. As our Base encoder is a custom implementation of the standard transformer encoder, it should be very close in implementation to Casanovo’s encoder, and thus the overall models are nearly identical. The disparity in the performance between our Base model and Casanovo’s reported result on nine-species, despite their similarities, cannot fully be explained in this work. One possible factor is that we were unable to identify the ideal hyperparameters for the model, due to the computational demands of the nine-species benchmark, which made it difficult to exhaustively optimize all settings. Furthermore, our focus was on evaluating pairwise features’ contribution to performance in isolation, thus we largely copied the hyperparameters chosen by Casanovo to make a fair comparison.

Uncertainty in the data and hyperparameters notwithstanding, we tested all models, including Casanovo’s provided checkpoint, on an independent bacterial dataset, applying our same model checkpoints as used in Table 3. With this dataset we found consistency between Base and PA, and now a considerable improvement over Casanovo. This evaluation further supports the merits of using pairwise features as input to the attention mechanism. Since pairwise features integrate seamlessly with transformer-based de novo models, with consistently higher performance in both our nine-species ablation studies and on our independent bacterial dataset, we believe this is an important contribution to the field as de novo sequencing becomes mainstream alongside traditional database searches.

## Acknowledgements

We kindly acknowledge the guidance of Wout Bittremieux and Melih Yilmaz regarding which specific versions of the nine-species datasets to use and where to acquire the data. We also acknowledge Daniela Klaproth-Andrade for her valuable input on earlier drafts of this manuscript. The computations were enabled by the Berzelius resource provided by the Knut and Alice Wallenberg Foundation at the National Supercomputer Centre in Sweden. LK and AN was supported by a grant from the Knut and Alice Wallenberg’s Foundation (KAW 2022.0032); Wallenberg AI, Autonomous Systems and Software Program (WASP) funded by the Knut and Alice Wallenberg Foundation; and by the Swedish Research Council (Grant 2024-05887). JL and MW were supported by an ERC Starting Grant (grant number 101077037).

## Footnotes

1 https://www.matrixscience.com/help/data_file_help.html

## References

1 Ruedi Aebersold and Matthias Mann. Mass-spectrometric exploration of proteome structure and function. Nature, 537(7620):347–355, September 2016.

2 Jürgen Cox and Matthias Mann. MaxQuant enables high peptide identification rates, individualized p.p.b.-range mass accuracies and proteome-wide protein quantification. Nature Biotechnology, 26(12):1367–1372, December 2008.

3 David N. Perkins, Darryl J. C. Pappin, David M. Creasy, and John S. Cottrell. Probability-based protein identification by searching sequence databases using mass spectrometry data. 20(18):3551–3567, 1999.

4 Viktoria Dorfer, Peter Pichler, Thomas Stranzl, Johannes Stadlmann, Thomas Taus, Stephan Winkler, and Karl Mechtler. MS Amanda, a Universal Identification Algorithm Optimized for High Accuracy Tandem Mass Spectra. Journal of Proteome Research, 13(8):3679–3684, August 2014.

5 Sangtae Kim and Pavel A. Pevzner. MS-GF+ makes progress towards a universal database search tool for proteomics. Nature Communications, 5(1):5277, October 2014.

6 Rui Qiao, Ngoc Hieu Tran, Lei Xin, Xin Chen, Ming Li, Baozhen Shan, and Ali Ghodsi. Computationally instrument-resolution-independent de novo peptide sequencing for high-resolution devices. Nature Machine Intelligence, 3(5):420–425, May 2021.

7 Melih Yilmaz, William E. Fondrie, Wout Bittremieux, Carlo F. Melendez, Rowan Nelson, Varun Ananth, Sewoong Oh, and William Stafford Noble. Sequence-to-sequence translation from mass spectra to peptides with a transformer model. Nature Communications, 15(1):6427, July 2024.

8 Tingpeng Yang, Tianze Ling, Boyan Sun, Zhendong Liang, Fan Xu, Xiansong Huang, Linhai Xie, Yonghong He, Leyuan Li, Fuchu He, Yu Wang, and Cheng Chang. Introducing π-HelixNovo for practical large-scale de novo peptide sequencing. Briefings in Bioinformatics, 25(2):bbae021, March 2024.

9 Ngoc Hieu Tran, Xianglilan Zhang, Lei Xin, Baozhen Shan, and Ming Li. De novo peptide sequencing by deep learning. Proceedings of the National Academy of Sciences of the United States of America, 114(31):8247–8252, August 2017.

10 Bin Ma. Novor: Real-Time Peptide de Novo Sequencing Software. Journal of The American Society for Mass Spectrometry, 26(11):1885–1894, November 2015.

11 Ngoc Hieu Tran, Rui Qiao, Zeping Mao, Shengying Pan, Qing Zhang, Wenting Li, Lei Xin, Ming Li, and Baozhen Shan. NovoBoard: a comprehensive framework for evaluating the false discovery rate and accuracy of de novo peptide sequencing, April 2024.

12 Douwe Schulte, Marta Šiborová, Lukas Käll, and Joost Snijder. Simultaneous polyclonal antibody sequencing and epitope mapping by cryo electron microscopy and mass spectrometry – a perspective. eLife, January 2025.

13 Sarah C. Jenson, Fanny Chu, Anthony S. Barente, Dustin L. Crockett, Natalie C. Lamar, Eric D. Merkley, and Kristin H. Jarman. MARLOWE: Taxonomic Characterization of Unknown Samples for Forensics Using De Novo Peptide Identification, September 2024.

14 Ashish Vaswani, Noam Shazeer, Niki Parmar, Jakob Uszkoreit, Llion Jones, Aidan N Gomez, L ukasz Kaiser, and Illia Polosukhin. Attention is all you need. In I. Guyon, U. Von Luxburg, S. Bengio, H. Wallach, R. Fergus, S. Vishwanathan, and R. Garnett, editors, Advances in Neural Information Processing Systems, volume 30. Curran Associates, Inc., 2017.

15 Wout Bittremieux, Varun Ananth, William E. Fondrie, Carlo Melendez, Marina Pominova, Justin Sanders, Bo Wen, Melih Yilmaz, and William S. Noble. Deep learning methods for de novo peptide sequencing. Mass Spectrometry Reviews, Nov 2024.

16 Terri Addona and Karl Clauser. De Novo Peptide Sequencing via Manual Interpretation of MS/MS Spectra. Current Protocols in Protein Science, 27(1):16.11.1–16.11.19, 2002.

17 Ragav Venkatesan and Baoxin Li. Convolutional Neural Networks in Visual Computing: A Concise Guide. CRC Press, Inc., 2017.

18 Siddharth Srivastava and Gaurav Sharma. OmniVec: Learning robust representations with cross modal sharing. In 2024 IEEE/CVF Winter Conference on Applications of Computer Vision (WACV), pages 1225–1237, Los Alamitos, CA, USA, January 2024. IEEE Computer Society.

19 Xuan Shen, Yaohua Wang, Ming Lin, Yilun Huang, Hao Tang, Xiuyu Sun, and Yanzhi Wang. DeepMAD: Mathematical Architecture Design for Deep Convolutional Neural Network. In 2023 IEEE/CVF Conference on Computer Vision and Pattern Recognition (CVPR), pages 6163–6173, Los Alamitos, CA, USA, June 2023. IEEE Computer Society.

20 Yanghao Li, Yuntao Chen, Naiyan Wang, and Zhao-Xiang Zhang. Scale-Aware Trident Networks for Object Detection. In 2019 IEEE/CVF International Conference on Computer Vision (ICCV), pages 6053–6062, Los Alamitos, CA, USA, November 2019. IEEE Computer Society.

21 Wenhai Wang, Jifeng Dai, Zhe Chen, Zhenhang Huang, Zhiqi Li, Xizhou Zhu, Xiaowei Hu, Tong Lu, Lewei Lu, Hongsheng Li, Xiaogang Wang, and Yu Qiao. Internimage: Exploring large-scale vision foundation models with deformable convolutions. In 2023 IEEE/CVF Conference on Computer Vision and Pattern Recognition (CVPR), pages 14408–14419, 2023.

22 John Jumper, Richard Evans, Alexander Pritzel, Tim Green, Michael Figurnov, Olaf Ronneberger, Kathryn Tunyasuvunakool, Russ Bates, Augustin Žídek, Anna Potapenko, Alex Bridgland, Clemens Meyer, Simon A. A. Kohl, Andrew J. Ballard, Andrew Cowie, Bernardino Romera-Paredes, Stanislav Nikolov, Rishub Jain, Jonas Adler, Trevor Back, Stig Petersen, David Reiman, Ellen Clancy, Michal Zielinski, Martin Steinegger, Michalina Pacholska, Tamas Berghammer, Sebastian Bodenstein, David Silver, Oriol Vinyals, Andrew W. Senior, Koray Kavukcuoglu, Pushmeet Kohli, and Demis Hassabis. Highly accurate protein structure prediction with AlphaFold. Nature, 596(7873):583–589, August 2021.

23 Mingxun Wang, Jian Wang, Jeremy Carver, Benjamin S. Pullman, Seong Won Cha, and Nuno Bandeira. Assembling the Community-Scale Discoverable Human Proteome. Cell Systems, 7(4):412–421.e5, October 2018.

24 Joon-Yong Lee, Hugh D Mitchell, Meagan C Burnet, Ruonan Wu, Sarah C Jenson, Eric D Merkley, Ernesto S Nakayasu, Carrie D Nicora, Janet K Jansson, Kristin E Burnum-Johnson, and Samuel H Payne. Uncovering hidden members and functions of the soil microbiome using DE novo metaproteomics. J. Proteome Res., 21(8):2023–2035, August 2022.

25 W. Fondrie and contributors. Depthcharge: A tool for peptide de novo sequencing, 2024.

26 Ngoc Hieu Tran, Xianglilan Zhang, Lei Xin, Baozhen Shan, and Ming Li. De novo peptide sequencing by deep learning. Proceedings of the National Academy of Sciences, 114(31):8247–8252, 2017.

