## Supplementary Figures for "Pairwise Attention: Leveraging Mass Differences to Enhance De Novo Sequencing of Mass Spectra"

### A Supplementary Figures

Table S1: **Average peptide precision (AUC) on V1 and V2 of the nine-species dataset**

| Test Set | Nine-species V1 | | | Nine-species V2 | | |
|---|---|---|---|---|---|---|
| Model | Base | PA | Casanovo | Base | PA | Casanovo |
| *Apis Mellifera* | 0.735 | **0.802** | 0.767 | 0.699 | 0.759 | **0.768** |
| *Bacillus Subtilis* | 0.852 | **0.899** | 0.866 | 0.799 | **0.861** | 0.832 |
| *Candidatus Endoloripes* | 0.700 | **0.750** | 0.720 | 0.671 | 0.742 | **0.772** |
| *Homo Sapiens* | 0.681 | **0.738** | 0.722 | 0.754 | 0.817 | **0.831** |
| *Methanosarcina Mazei* | 0.852 | **0.899** | 0.866 | 0.799 | **0.861** | 0.832 |
| *Mus Musculus* | 0.770 | **0.806** | 0.766 | 0.700 | **0.747** | 0.721 |
| *Solanum Lycopersicum* | 0.833 | **0.890** | 0.837 | 0.831 | 0.881 | **0.883** |
| *Saccharomyces Cerevisiae* | 0.855 | **0.889** | 0.871 | 0.833 | **0.879** | 0.850 |
| *Vigna Mungo* | 0.868 | **0.892** | 0.849 | 0.811 | **0.870** | 0.746 |
| Average | 0.791 | **0.836** | 0.804 | 0.767 | **0.824** | 0.804 |

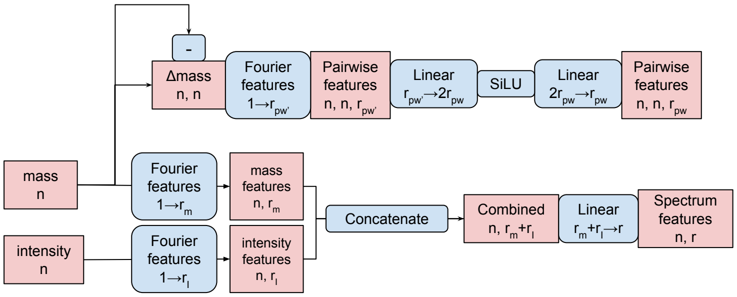


**(a)**

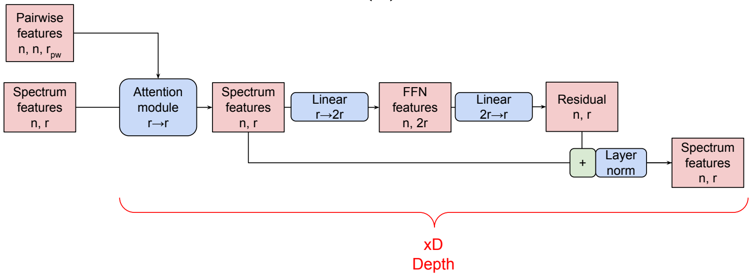


**(b)**

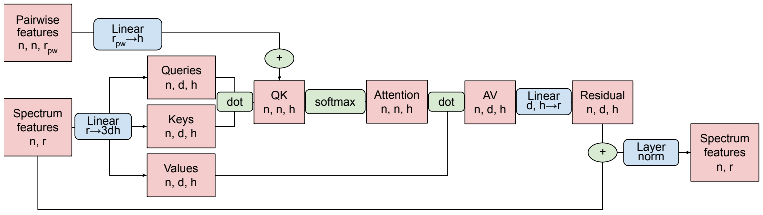


**(c)**

Figure S1: Schematic depicting the architectural integration of pairwise features into the Pairwise Attention model's encoder. In a) the preprocessing of mass and intensity vectors produces both spectral and pairwise features, which are then input into the main transformer blocks. In b) the repetitive unit of the encoder is outlined. Spectral features are fed into the first block of the encoder, while pairwise features are fed into every attention module throughout the depth of the encoder. The mechanism of communication between the spectral features and pairwise features is depicted in c). Besides the pairwise features, this is a standard self-attention module, but with them, there is an extra linear transformation to the number of heads and an addition operation as a bias before the softmax. Not pictured is the transformer decoder, which for this project is the native pytorch layer implementation.

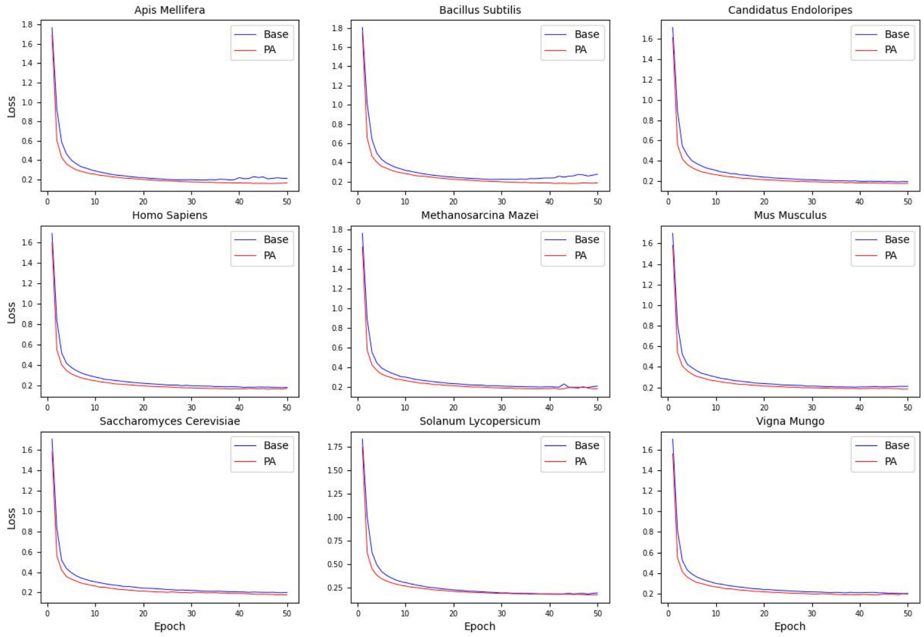


Figure S2: Training loss as a function of epoch for V1 of the nine-species dataset for our base and pairwise models. Training loss was averaged over every epoch.

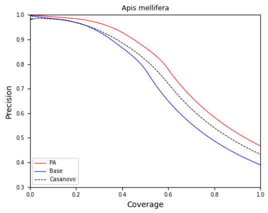


(a) *Apis mellifera*

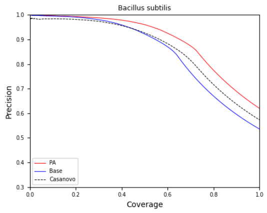


(b) *Bacillus subtilis*

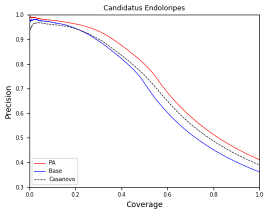


(c) *Candidatus endoloripes*

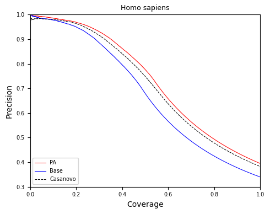


(d) *Homo sapiens*

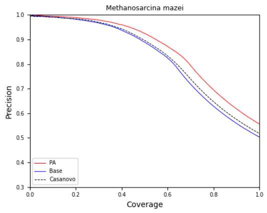


(e) *Methanosarcina mazei*

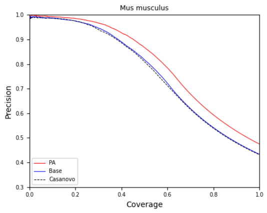


(f) *Mus musculus*

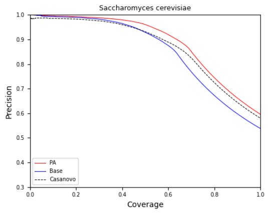


(g) *Saccharomyces cerevisiae*

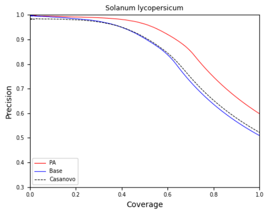


(h) *Solanum lycopersicum*

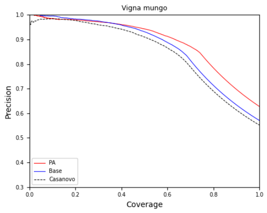


(i) *Vigna mungo*

Figure S3: **Peptide precision vs. coverage on the nine-species V1 dataset, training on 8 species and evaluating on the held-out 9th.** All individual species are displayed.

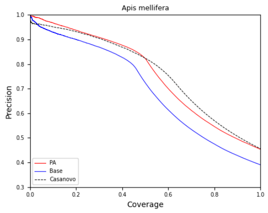


(a) *Apis mellifera*

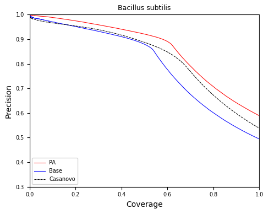


(b) *Bacillus subtilis*

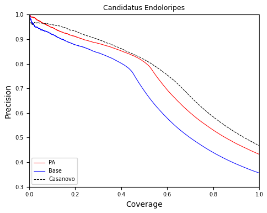


(c) *Candidatus endoloripes*

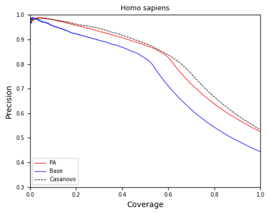


(d) *Homo sapiens*

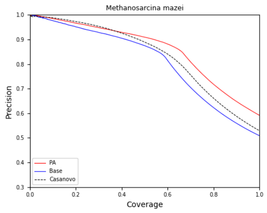


(e) *Methanosarcina mazei*

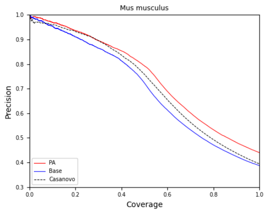


(f) *Mus musculus*

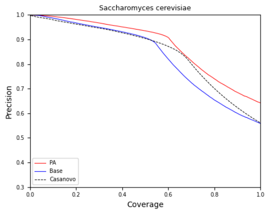


(g) *Saccharomyces cerevisiae*

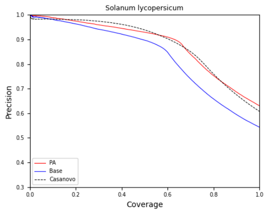


(h) *Solanum lycopersicum*

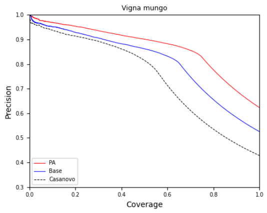


(i) *Vigna mungo*

Figure S4: **Peptide precision vs. coverage on the nine-species V2 dataset, training on 8 species and evaluating on the held-out 9th.** All individual species are displayed.

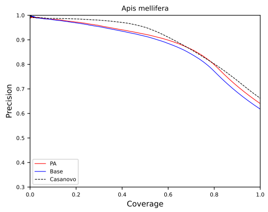


(a) *Apis mellifera*

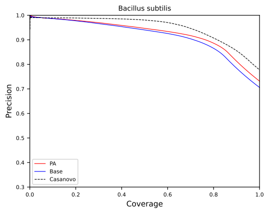


(b) *Bacillus subtilis*

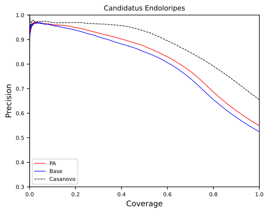


(c) *Candidatus endoloripes*

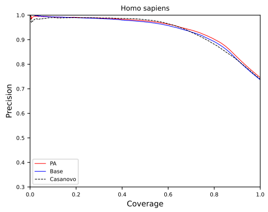


(d) *Homo sapiens*

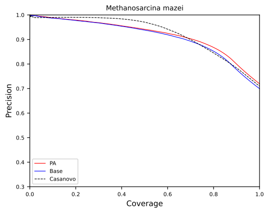


(e) *Methanosarcina mazei*

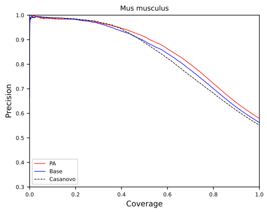


(f) *Mus musculus*

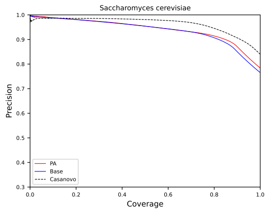


(g) *Saccharomyces cerevisiae*

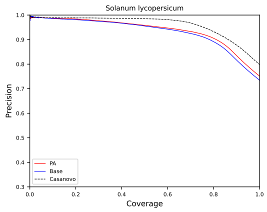


(h) *Solanum lycopersicum*

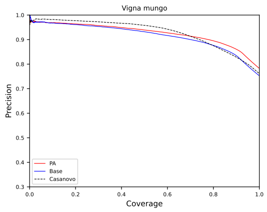


(i) *Vigna mungo*

Figure S5: **Peptide precision vs. coverage on the nine-species V2 dataset after training on the MassIVE-KB dataset.** All individual species results are shown separately.
